# Liposome-based transfection enhances RNAi and CRISPR-mediated mutagenesis in non-model nematode systems

**DOI:** 10.1101/429126

**Authors:** Sally Adams, Prachi Pathak, Hongguang Shao, James B. Lok, Andre Pires-daSilva

## Abstract

Nematodes belong to one of the most diverse animal phyla. However, functional genomic studies in nematodes, other than in a few species, have often been limited in their reliability and success. Here we report that by combining liposome-based technology with microinjection, we were able to establish a wide range of genomic techniques in the newly described nematode genus *Auanema*. The method also allowed heritable changes in dauer larvae of *Auanema*, despite the immaturity of the gonad at the time of the microinjection. As proof of concept for potential functional studies in other nematode species, we also induced RNAi in the free-living nematode *Pristionchus pacificus* and targeted the human parasite *Strongyloides stercoralis*.

## Introduction

Traditionally, the analysis of gene function has been restricted to a few model systems because techniques for manipulating gene expression in one species are often not easily transferable to other species. However, recent technological advances in high-throughput sequencing, gene silencing strategies, and genome editing have changed this and opened the door to mechanistic studies in non-model systems. This allows us to ask questions that cannot be addressed with well-established genetic model organisms^1,2^.

Nematodes are one of the most diverse animal phyla, with an estimated 40 million species^3^. *Caenorhabditis elegans* is the best-studied nematode species and has been used as a model for research in developmental genetics, cell biology, and evolution^4^. *Auanema rhodensis* is also a member of the Eurhabditis clade^5^, and its size^6^, ecology^7^ and culture biology are similar to *C. elegans* ^8^. *A. rhodensis’* unique features make it an interesting model for comparative studies^9^. For example, instead of the usual pattern of X chromosome inheritance from mother-to-son, *A. rhodensis* sons inherit the X chromosome from the father^10^. Moreover, *A. rhodensis* males produce haploid cells that are discarded as female polar bodies^11,12^, and sex determination is influenced by the mother’s age^13^. *A. rhodensis* is unique among hermaphroditic nematodes in that two distinct groups of stem cells in the hermaphrodite germline produce either sperm or oocytes^14^. Thus, *Auanema* is a suitable model for basic cell biology, genetics, and evolution. However, the development of tools for gene manipulation is a crucial requirement for this.

One of the most broadly applicable tools is RNA interference (RNAi), which is a conserved biological response to double-stranded RNA (dsRNA). RNAi inactivates gene expression in a sequence-specific manner^15^. In some nematodes, delivery of RNAi triggers can be performed simply by soaking them in a dsRNA solution, circumventing the need for microinjection^16–20^. However, this method is often inefficient or fails entirely in many nematodes. They might lack effector genes that are either required for the uptake of the dsRNA (environmental RNAi response) and/or the amplification of the RNAi effect throughout the animal and on to the next generation (systemic RNAi response)^21–25^. In a range of nematodes the RNAi response can be improved via the addition of chemicals (e.g., serotonin, octopamine, spermidine, lipofectamine) that enhance uptake of the dsRNA/siRNA from the medium^26–30^. Another option is the direct injection of dsRNA into the body cavity or gonad^22,31,32^.

However, even with these modifications, the efficiency of transcriptional-silencing can vary dramatically between targets. In some cases, the greatest efficiency is observed in the tissues most accessible to the soaking solution^29^. In *C. brenneri*, injection of dsRNA results in a phenotype only if the dsRNA is directly injected into the gonad and not into the body cavity, suggesting that physical barriers can modulate the RNAi response^17,22^. Furthermore, many species are resistant to RNAi, even by direct injection of the dsRNA^22,33,34^. Although this might be due to a lack of effector genes, differences in physiology or morphology of the gonad may also play a role. The addition of a substance that effectively spreads RNA from the site of injection can help to overcome the limitations of a less robust spreading system.

In our initial attempts to establish an RNAi protocol for *Auanema*, we employed conventional methods. However, soaking, electroporation or microinjection of dsRNA were unsuccessful. One possible reason could be differences in the gonad morphology between *Auanema* and *C. elegans*. In *C. elegans* hermaphrodites, the distal gonad arms contain germline nuclei within a common cytoplasm, forming a syncytium. This facilitates microinjection because the injected material can be incorporated by many oocytes as they mature and become cellularised, eventually being incorporated into the fertilized egg^35^. It is not clear that the germline in *Auanema* has a syncytium, given the very thin rachis^14^. Consequently, when nucleic acids are introduced into the gonad by microinjection, it is probable that they reach only a limited number of germline cells. In addition, the mitotic region of the *Auanema* gonad (the usual target for microinjection) is much shorter and more difficult to identify than in *C. elegans*^6^ (Figure 1).

**Figure 1.**
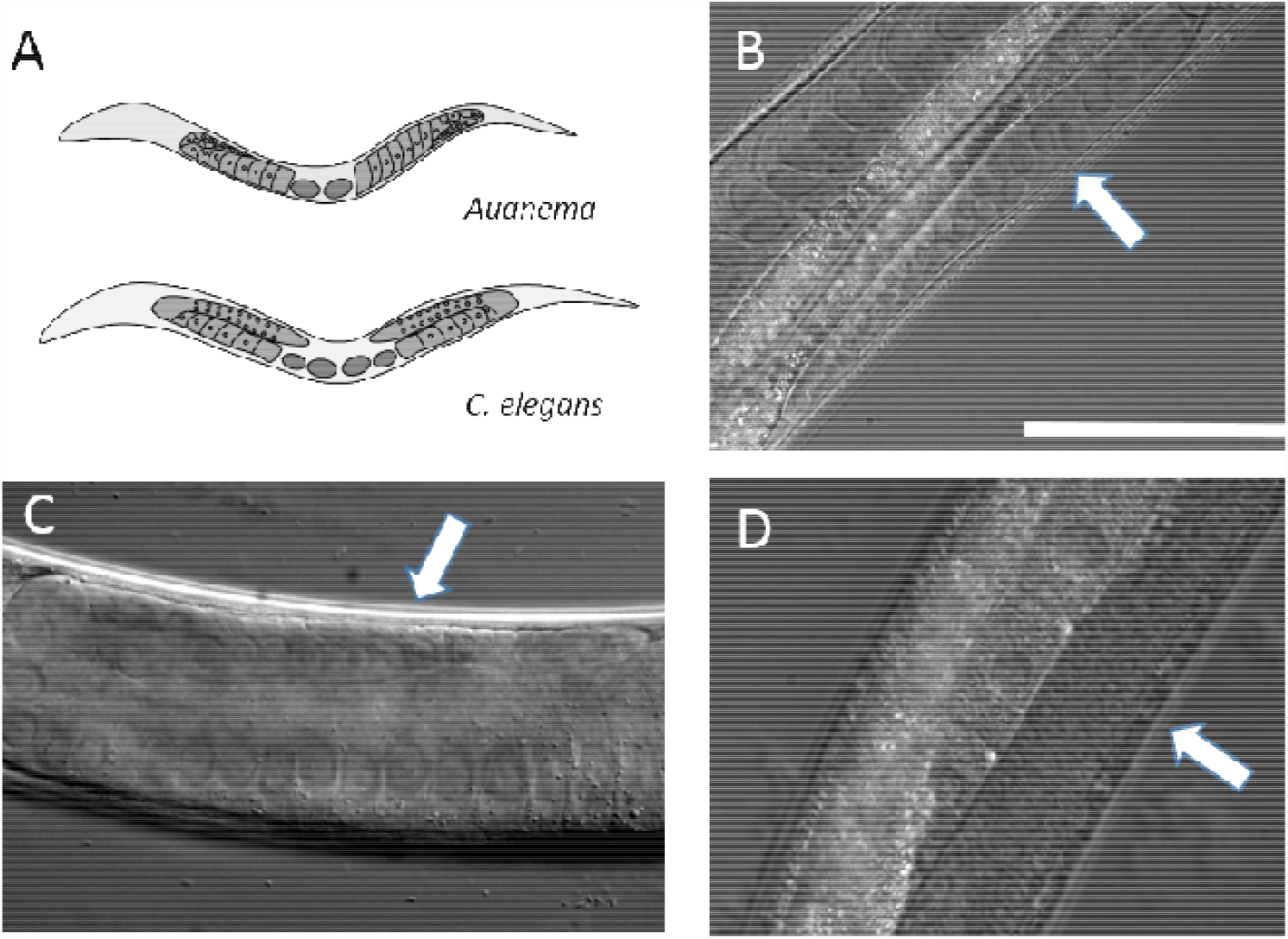
Morphological differences between the gonads of the genus *Auanema* and *C. elegans* (A). Schematic diagram illustrating the arrangement of the gonad arms in *Auanema* (top) and *C. elegans* (bottom). The distal gonad arm of *A. rhodensis*(B) and *A. freiburgensis*(C) is smaller, contains fewer, larger nuclei compared *C. elegans* (D). The white arrow highlights the distal arm. Scale bar is 50 µM.

Liposomes have been used as transfection agents to deliver various materials to cells. Liposomes consist of vesicles, bounded by a lipid bilayer, that can be loaded with cargo such as nucleic acids and/or proteins^36–39^. The transfection reagent lipofectamine has been used previously in nematodes to improve siRNA uptake from the environment^29^. However, this protocol also failed in *A. rhodensis*, prompting us to try it in combination with microinjection.

Here we report that by combining liposome-based technology with microinjection into the gonad, functional genomic techniques become feasible in non-model nematode species. With this method, RNAi and CRISPR-Cas9 mutagenesis become extremely efficient in *A. rhodensis* and *A. freiburgensis*, often inducing a phenotype in the entire brood of the injected animal. We also often observed RNAi-or CRISPR-Cas9-induced phenotypes in the injected (P0) nematode, suggesting that the liposomes spread from the gonad to somatic cells. We were also able to transfect developmentally arrested larval stages, such as the dauer larvae of *A. rhodensis*, and the infective third larval stage (iL3) of the mammalian parasitic nematode *Strongyloides stercoralis*. In addition, we established RNAi in *Pristionchus pacificus*, a satellite model system to *C. elegans*, that is used for comparative studies. Thus, this method makes a widening range of nematodes amenable to functional studies.

## Materials and methods

### Nematodes strains and culture

We used inbred strains of the free-living nematodes *Auanema rhodensis* APS4, *A. freiburgensis* APS7^10^, and *Pristionchus pacificus* PS312 derivative “97”^40^. These strains were maintained in the laboratory on nematode growth medium (NGM) agar plates seeded with the streptomycin-resistant *Escherichia coli* OP50–1 strain and cultured at 20 °C by standard procedures for *C. elegans*^41^. Microbial contamination was prevented by adding 200 µg/mL nystatin and 200 µg/mL streptomycin to the NGM ^42^.

We used the UPD strain of the parasitic nematode *S. stercoralis*, which was originally isolated from naturally infected dogs in 1976. Free-living adults and iL3 were obtained from charcoal coprocultures from immunosuppressed infected dogs^43,44^. *S. stercoralis* was maintained in adult mixed-breed dogs that were bred and housed in a vivarium at the University of Pennsylvania that is accredited by the US Association for the Assessment and Accreditation of Laboratory Animal Care. The vivarium was climate controlled, and the dogs were fed on standard laboratory chow and given regular husbandry and medical care by the technical and veterinary staff of the University Laboratory Animal Resources (ULAR) group at the University of Pennsylvania. All maintenance of dogs and related procedures were conducted in accordance with protocols 804798 and 804883 (for parasite maintenance) and 803357 (for dog breeding) approved by the University of Pennsylvania Institutional Animal Care and Use Committee (IACUC). All IACUC protocols and routine husbandry and medical care of animals at the University of Pennsylvania were conducted in strict accordance with the Guide for the Care and Use of Laboratory Animals of the US National Institutes of Health ^45^.

### Production of dsRNA for RNAi

Homologues of *par-1* and *unc-22* were identified by BLAST searches of the *A. rhodensis* genome or the *A. freiburgensis* transcriptome with the *C. elegans* proteins PAR-1 (H39E23.1a) and UNC-22 (Q23551) as queries (genome and transcriptome to be published elsewhere)^46^. *Arh-par-1*(MH249768), *Afr-par-1* (MH249770), *Arh-unc-22* (MH249769) and *Afr-unc-22* (MH124555) encode predicted proteins of 1051, 637, 6735 and 4095 amino acids, respectively. *Arh*-PAR-1 and *Afr*-PAR-1 share 49 and 48% identity, respectively, with the *Cel*-PAR-1 (Figure S1a). It is probable that *Afr-par-1* is a partial sequence that lacks the C-terminal encoding region. *Arh-unc-22* and *Afr-unc-22* encode proteins that share 69% and 74% sequence identity, respectively, with *Cel*-UNC-22. *Afr-unc-22* cDNA is also likely to be a partial sequence, lacking the start of the coding region.

To produce target-specific dsRNA, we used a gene-specific PCR product as template. For *par-1* RNAi, PCR primers were designed to amplify a 958 bp (*Arh-par-1*), 689 bp (*Afr-par-1*) or 756 bp (*Ppa-par-1*, PPA12901) fragment from a sequence encoding the predicted N-terminal region, overlapping the conserved kinase domain (Figure S1b, see Table S1 for primer sequences). *unc-22* RNAi PCR primers were designed to span 700 bp of *Arh-unc-22* and 1,516 bp of the *Afr-unc-22* cDNA sequence (Figure S2, Supplemental Table 1). For *in vitro* transcription of both strands, the T7 promoter sequence (5’-TAATACGACTCACTATAGGG-3’) was included at the 5’ end of the primer sequences. For the synthesis of cDNA, RNA was isolated from mixed stage cultures. In brief, nematodes were washed from a mixed stage culture on a six cm plate with 500 µl of M9 buffer^41^. Nematodes were allowed to settle by gravity sedimentation and the supernatant was removed. The pellet was then washed two more times with M9 buffer using gravity sedimentation. After the final wash, the supernatant was removed; 200 µl of Trizol (Ambion) was added, and the sample stored at –80 °C. To extract RNA, each sample was subjected to 3 rounds of freeze-cracking, alternating between freezing on dry ice and thawing at room temperature. After the final freeze-cracking step, another 200 µl of Trizol^®^ were added to each tube and RNA was isolated according to the manufacturer’s protocol. cDNA was synthesised from 0.5 µg of total RNA using M-MLV reverse transcriptase (Promega) and random primers (Promega) according to the manufacturer’s instructions. The required cDNA fragment was then amplified using gene-specific T7 promoter primers. A 20 µl reaction was carried out with GoTaq© Green Master Mix (Promega), with approximately 200 ng of cDNA and 250 nM of each primer using 30 cycles at the following conditions: 94 °C for 15 s, 52 °C for 30 s and 72 °C for 30 s. The amplified fragment was cleaned using the QIAquick PCR Purification kit (Qiagen), according to the manufacturer’s protocol, and eluted in 30 µl of elution buffer. dsRNA was then transcribed *in vitro* at 37 °C for 4 h from approximately 200 ng of the PCR template using T7 RNA polymerase (Promega), following the manufacturer’s instructions. *In vitro* transcripts were cleaned with the RNA clean-up protocol in the RNeasy^®^ Mini Kit (Qiagen). The microinjection mixture was produced by incubating 50 ng/µl of dsRNA and 12% Lipofectamine^®^RNAiMax reagent (Invitrogen) in a 10 µl total volume of nuclease-free water at room temperature for 20 min. For non-lipofectamine controls, the injection mixture was prepared as above but Lipofectamine^®^RNAiMax reagent was omitted and its volume substituted with additional RNase-free water. The mixture was then used immediately for microinjection (see details below).

### CRISPR-Cas9 directed mutagenesis

Mutations in the collagen genes *Cel-rol-6*(*su1006*) and *Ppa-prl-1*(*tu92*), which convert an arginine residue into cysteine, result in a dominant roller phenotype^47,48^. To determine if the same mutation could be used as a dominant marker in other species, we used the *C. elegans* ROL-6 protein (P20784) to identify candidate sequences in the transcriptome of *A. freiburgensis* and the genomes of *A. rhodensis* and *S. stercoralis*. In each case, multiple sequences were obtained that encode ROL-6-like collagen proteins (Figures S3, S4). *Afr-rol-6.1* (MH124551), *Arh-rol-6.1*(MH124552) and *S. stercoralis*(SSTP_0000742500) encode the proteins that share the highest level of sequence identity with *Cel*-ROL-6 (63, 62 and 44%, respectively), compared to other candidates (Figures 3A, S3, S4). To obtain the genomic DNA sequence of *Afr-rol-6.1* (as no genome is currently available), we used the primers UW174 and UW175 (Supplemental table S1) to amplify a genomic region flanking the target site, employing the single-nematode PCR method^49^. The resulting amplicon was sequenced with the primers used for PCR amplification and used as a template to design CRISPR guide RNA (crRNA) and donor fragments as described below (Figure S5, S6).

The *Streptococcus pyogenes* Cas9 endonuclease requires two short RNAs, a sequence-specific CRISPR RNA (crRNA) and a universal transactivating RNA (tracrRNA). When combined, these RNAs hybridize through short complementary regions and form the small guide RNA (sgRNA), a complex that guides the Cas9 to the target DNA sequence. Site-specific recognition and cleavage by the endonuclease also require a protospacer adjacent motif (PAM) (5’-NGG-3’). This sequence is found immediately adjacent to the complementary DNA sequence and induces a conformational change in Cas9 that triggers the double-stranded break^50^.

To introduce the nucleotide substitutions required to change the codon from arginine to cysteine in each of the three subject nematodes, we used the oligonucleotide-mediated gene conversion strategy^51,52^ in combination with the Alt-R^™^ CRISPR-Cas9 System from Integrated DNA Technologies (IDT). With this strategy, double-stranded breaks are repaired by homologous recombination (HR) using a single-stranded DNA (ssDNA) oligonucleotide as a donor fragment provided in *trans*, allowing endogenous gene editing.

For each gene edit, a pair of antiparallel sgRNA were generated by annealing two gene-specific crRNAs and a tracrRNA (Figures 3B, S6, Supplemental Table 2). crRNAs and tracrRNA were synthesised by IDT. The sgRNA mixes were prepared as follows: crRNAs and tracrRNA were resuspended in nuclease-free water to a stock concentration of 100 µM; 1.5 µl of each crRNA and 3 µl of tracrRNA from the prepared stocks were added to 4 µl duplex buffer (provided by IDT) to an end volume of 10 µl of mixed RNA solution (30 µM final concentration). To generate the anti-parallel sgRNA, the mixed RNA solution was incubated at 95 °C for 5 minutes, followed by cooling to room temperature. In our hands, the expected roller phenotype was not produced if either of the crRNAs was used individually (data not shown). The gene-specific sequences of all sgRNA are listed in Supplemental Table 2. For each gene edit, a ssDNA PAGE-purified oligonucleotide of 100 bases that incorporates the required nucleotide changes was synthesised commercially (IDT) and used as donor fragment (Figure 3b, S6, Supplemental table 2). Injection mixtures for CRISPR-Cas9 designed to mediate gene conversion in both *Auanema* spp. and *S. stercoralis* were prepared as follows: ribonucleoprotein (RNP) complexes were assembled by incubating 25 µg (2.5 µl) of Alt-R^™^ *S. pyogenes* Cas9 Nuclease 3NLS (IDT) with 2.5 µl of the 30 µM sgRNA stock solution at room temperature for 5 min and 1 µl of ssDNA donor fragment (200 ng/µl) was then added to a final concentration of 33 ng/µl to give a final volume of 6 µl. To produce the liposome-based microinjection mixes for both *Auanema* spp. and *S. stercoralis*, we diluted the RNP-ssDNA mixture 5-fold in RNase-free M9 buffer, added 3% Lipofectamine^®^RNAiMax reagent (v/v) (Thermofisher Scientific) and incubated at room temperature for 20 min. The mixture was then immediately microinjected as described below.

### Construction of pAS19 (P*eef-1A.1*::TurboRFP::*tbb-2*-3’UTR)

To develop transgene expression protocols for *Auanema*, we built the expression construct pAS19, which incorporates an *A. rhodensis-* specific promoter and 3’ untranslated region (3’UTR) flanking a TurboRFP reporter gene (Figure S12)^48^. The *eef-1A.1* promoter was chosen, as it drives robust germline expression in *C. elegans* and has proven invaluable in CRISPR-Cas9 techniques^53^. *Arh-eef-1A.1* (MH124553) was identified using *C. elegans* EEF-1A.1 protein (P53013) as the query in a local BLAST search of the *A. rhodensis* genome. The predicted *A. rhodensis* protein resulting from this search exhibits 94% identity with *Cel*-EEF-1A.1 (Figure S8). A 926 bp region directly upstream of the predicted *Arh-eef-1A* start codon was synthesised as a gBlock gene fragment (IDT) (Figure S9). This fragment incorporates regions at the 5’ and 3’ ends that correspond to DNA sequences flanking the *Kpn1* site directly upstream of the turbo RFP coding sequence in *Ppa*::RFP^48^. Following its linearisation by digestion with *KpnI* (Promega), the gBlock gene fragment was inserted into *Ppa::* RFP by Gibson cloning according to the manufacturer’s instructions (NEB).

To further optimise expression in *A. rhodensis*, the *Cel-rpl-23* 3’ UTR was replaced by a predicted *Arh-tbb-2*3’ UTR. A BLAST search of the *A. rhodensis* genome with *Cel*-TBB-2 (P52275) identified *Arh-tbb-2*(MH124554), which encodes a predicted protein sharing 93% identity with its *C. elegans* homologue (Figure S10). A gBlock gene fragment (IDT) was synthesised, which incorporated a 238 bp fragment immediately downstream of the predicted stop codon of *Arh-tbb-2* (Figure S11). The *Cel-rpl-23* 3’ UTR was removed by digestion with *SacI* and *EcoRI* and replaced with the *Arh-tbb-2*3’UTR by Gibson cloning, according to the manufacturer’s instructions (NEB). For injection, 500 ng/µl of pAS19 were incubated for 20 min at room temperature with 12% Lipofectamine^®^RNAiMax reagent (v/v) (Thermofisher Scientific) and used immediately.

### Microinjection of *Auanema* and *P. pacificus*

Microinjections of *A. rhodensis, A. freiburgensis* and *P. pacificus* were conducted using an IM-300 Pneumatic Microinjector system with an oil hydraulic Micromanipulator (Narishige). Agarose pads for microinjection were prepared by adding a drop of molten 2% agarose (w/v) in water to the centre of 24 × 60 mm coverslips (Menzel-Glaser). Then a second coverslip was added quickly on top to produce a thin agarose layer. Once solidified, the top coverslip was removed and the agarose pad dried on a heating block for 1–2 h at 60 °C. 3 µl of the required injection mixture were loaded into pre-pulled microcapillary needles (Tritech Research) with an outside diameter of 1.0 mm, an inside diameter of 0.6 mm and a tip diameter of less than 1 µm. To immobilise nematodes for injection, young adults or dauer larvae were moved to a small drop of undiluted halocarbon oil 700 (Sigma) on the dried agarose pad and gently pressed onto it using either a platinum pick or eyelash. Adults were injected towards the distal end of each gonad, as close to the tip as possible, using an input gas pressure of 75 psi, balance pressure of 1.3 psi and an injection pressure of 20 psi. The injection of dauer larvae was more technically challenging and we found input gas pressure to be critical. To optimise conditions, we conducted a set of preliminary experiments, with a gradient of input gas pressures ranging from 50 to 75 psi and an increment of 5 psi each time. Dauer larvae injected with an input pressure of 70 psi or above did not survive, whilst those injected with input pressures below 60 psi never exhibited the roller phenotype. Therefore, all further injections of dauer larvae were conducted with an input gas pressure between 60 and 65 psi. After injection, all individuals were released from the agarose pad by adding a small quantity of M9 buffer^41^. They were then transferred to standard NGM plates seeded with OP50–1, for recovery and further analysis.

### Microinjection of *S. stercoralis*

The prepared mixes for CRISPR-Cas9-mediated gene conversion were delivered to free-living female *S. stercoralis* by microinjection into the syncytial region of the ovary, midway between the distal tip and the bend, using techniques adapted from *C. elegans* methodology^35,54,55^. Free-living females were immobilized for injection under drops of halocarbon oil on dry agar pads made from molten 2% agarose, as described above for *A. rhodensis* and *A. freiburgensis*. Glass microinjection needles with inner, outer and tip diameters, as described for *A. rhodensis* and *A. freiburgensis*, were prepared by finely drawing borosilicate glass capillary tubing with a vertical pipette puller (Model P30, Sutter Instrument, Novato, California, USA). Mounted worms were viewed with an inverted microscope (Model IX70, Olympus USA) with differential interference (DIC) optics at 400X magnification. Needles containing microinjection mixes were positioned with a micromanipulator (Leitz Wetzlar) and pressurized with nitrogen using a Pneumatic PicoPump microinjector (Model PV80, World Precision Instruments, Sarasota, Florida USA). Injection pressures for free-living female *S. stercoralis* were in the range of 30–50 psi. Parasitic nematode iL3s, including those of *S. stercoralis*, are radially constricted in configuration, and their cuticles are thickened and resistant to chemical and physical degradation compared to those of free-living forms. iL3 are generally more highly motile than their continuously developing counterparts. These morphological and behavioural characteristics necessitated some modifications in microinjection technique for *S. stercoralis* iL3. Thicker dry agar pads, made from molten 3.5% agarose, were necessary to immobilize iL3 for injection. Also, the internal hydrostatic pressure of iL3 appeared to be higher than in free-living female *S. stercoralis*, causing prolapse of internal contents from the worms if normal injection pressures were used. For this reason, injection pressures for iL3 were maintained at or below 20 psi. Positioning of the needle relative to the body of the iL3 was also crucial for the successful injection of *S. stercoralis* at this stage. The point of penetration was near the junction of the pharynx and intestine, and the needle was positioned at a 15° angle relative to the long axis of the worm. Care was taken to penetrate the cuticle of the iL3 as gently as possible, delivering injection mix into the pseudocoelom, while avoiding contact between the needle tip and the intestine. Both iL3 and free-living female *S. stercoralis* were demounted immediately after injection by gentle prodding with a “worm pick” fashioned from 36 gauge platinum wire. Then they were transferred to drops of water in the centres of standard 60 mm NGM plates with lawns of *E. coli* OP50^41^. Free-living females and iL3 injected by these methods generally recovered within 10–15 minutes of demounting.

### Genomic DNA extraction and genotyping of roller lines

Genomic DNA was extracted from approximately 500 mixed stage worms using the Gentra Puregene Tissue Kit (Qiagen), according to the manufacturer’s instructions but with the following modification. Collagenase I (Sigma) was added to a concentration of 0.2 % (w/v), after proteinase K digestion, and samples incubated at 37°C for 30 minutes. The *Arh-rol-6.1* locus was amplified with the primers UW159 and UW555 (See Supplemental Table 1) and Q5 high fidelity DNA polymerase (NEB), including the high GC enhancer, according to the manufacturer’s instructions. Amplicons were cleaned using the QIAquick PCR purification kit (Qiagen) and submitted for Sanger sequencing with primer UW555.

**Table 1.**
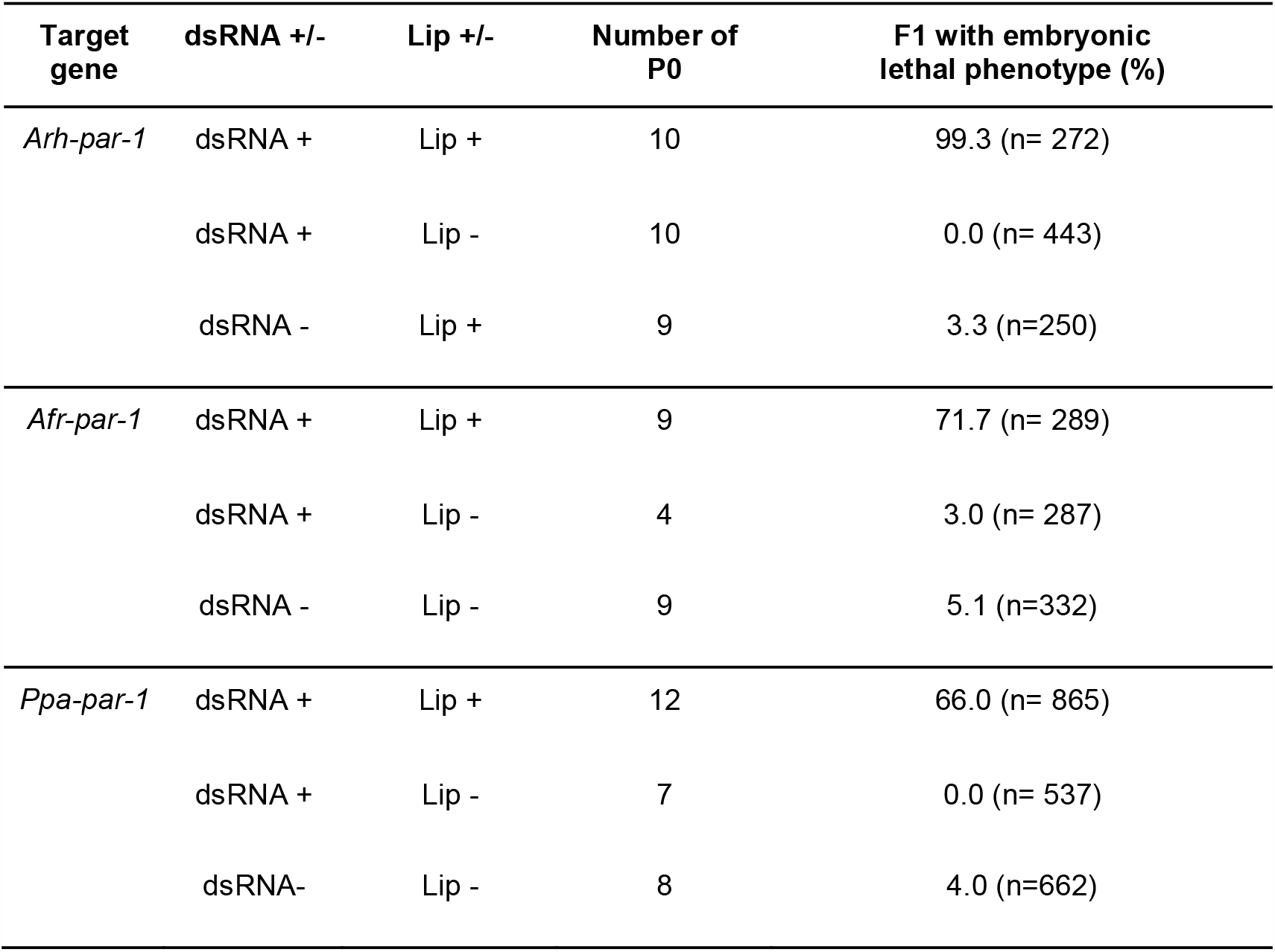
Summary of RNAi targeting *par-1* homologues in *A. rhodensis, A. freiburgensis*,and *P. pacificus*. Results shown are the combined data from at least 2 independent sets of injections.

## Results

### Lipofection-based methods dramatically improve RNAi efficiency in *A. rhodensis, A. freiburgensis*, and *P. pacificus*

To determine if liposome-based transfection could facilitate RNAi in *A. rhodensis* and *A. freiburgensis*, we targeted homologues of the *C. elegans* genes, *Cel-par-1* and *Cel-unc-22*. Both genes exhibit clear, easy-to-score phenotypes when targeted by RNAi in *C. elegans* ^15,56^. *Cel-par-1* encodes a maternally provided serine-threonine protein kinase that is required for establishing early embryonic polarity^57^ and RNAi by injection of *par-1* dsRNA results in embryonic lethality in 100% of the progeny^56^. The *Cel-unc-22* gene encodes twitchin, an abundant myofilament protein^58,59^. Down-regulation of the *unc-22* transcript by RNAi results in a severe twitching phenotype in the progeny of *C. elegans* injected with dsRNA^15^.

Injection of dsRNA targeting *Arh-par-1* or *Afr-par-1* resulted in high levels of embryonic lethality, but only if lipofectamine was included in the injection mixture (Table 1). By contrast, injection of only lipofectamine (12% v/v in water) did not result in significant embryonic lethality (Table 1), indicating that the observed phenotype results from down-regulation of the target and not from toxicity of the lipofectamine. To test the application of the method to other nematode species that have been resistant to conventional RNAi methods, we targeted the *Ppa-par-1* gene in *P. pacificus* (Table 1, Figure S1). As with both *Auanema* species, injection of *Ppa-par-1* dsRNA combined with lipofectamine resulted in high levels of embryonic lethality in the F1 (Table 1). Injection of *Ppa-par-1* dsRNA or lipofectamine (12% v/v in water) alone resulted in no embryonic arrest in the F1 (Table 1). To further test the liposome-based RNAi method, we targeted *unc-22* homologues in both *Auanema* species. Strikingly, the majority of the nematodes injected with dsRNA-liposome complexes (P0) exhibited a severe twitching phenotype 16 to 24 hours after injection, whereas no phenotype was observed with dsRNA alone (Table 2, Figure 2). The individuals that scored as ‘twitching’ not only exhibited uncoordinated muscle spasms but also had severely impaired movement. Individuals eventually became immobile and were only able to eat the bacteria in the site they occupied, resulting in easily identified cleared patches in the bacterial lawn around the heads of affected worms. Furthermore, in both species, this severe twitching phenotype was inherited by 100% of the F1 progeny of the P0 worms expressing the twitching phenotype (Table 2).

**Table 2.**
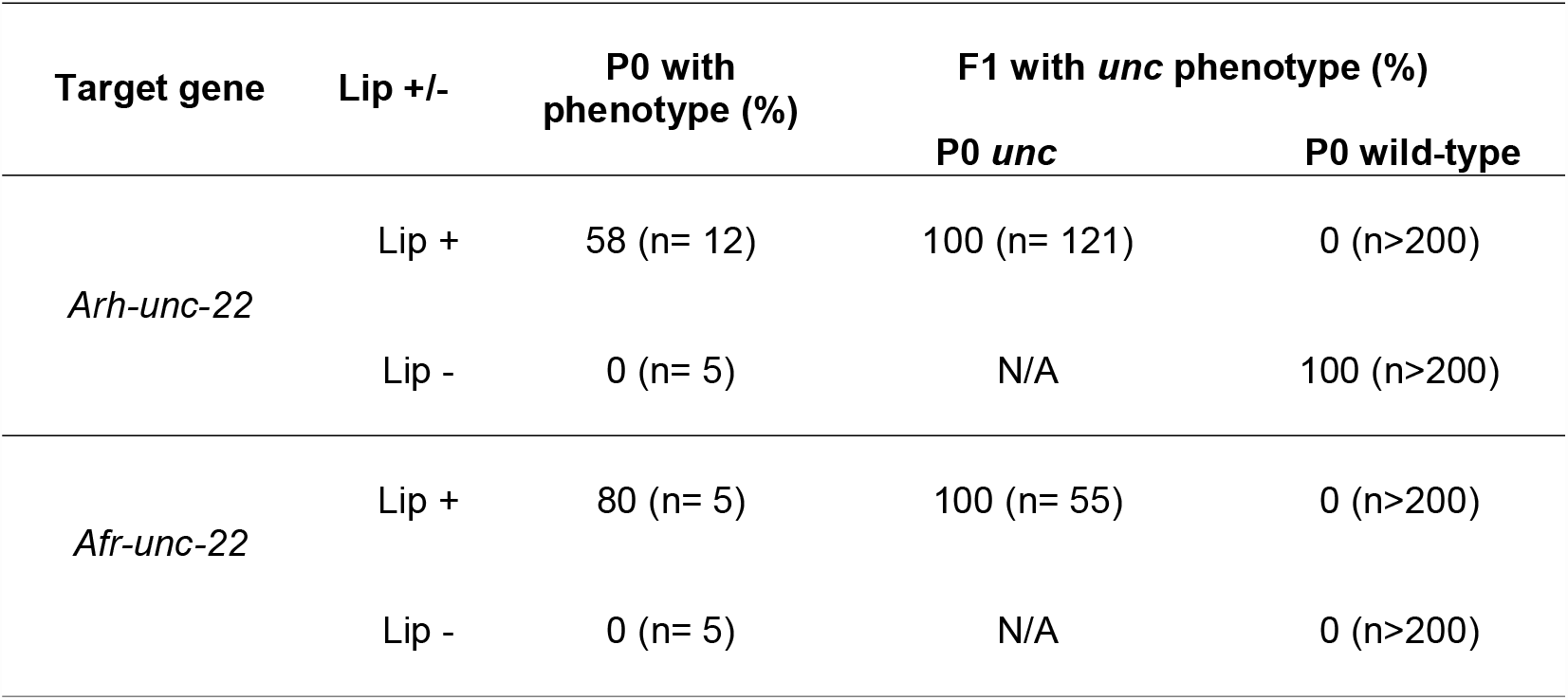
Summary of RNAi targeting *unc-22* homologues in *A. rhodensis* and *A. freiburgensis*. Results shown are the combined data from at least 2 independent sets of injections.

**Figure 2.**
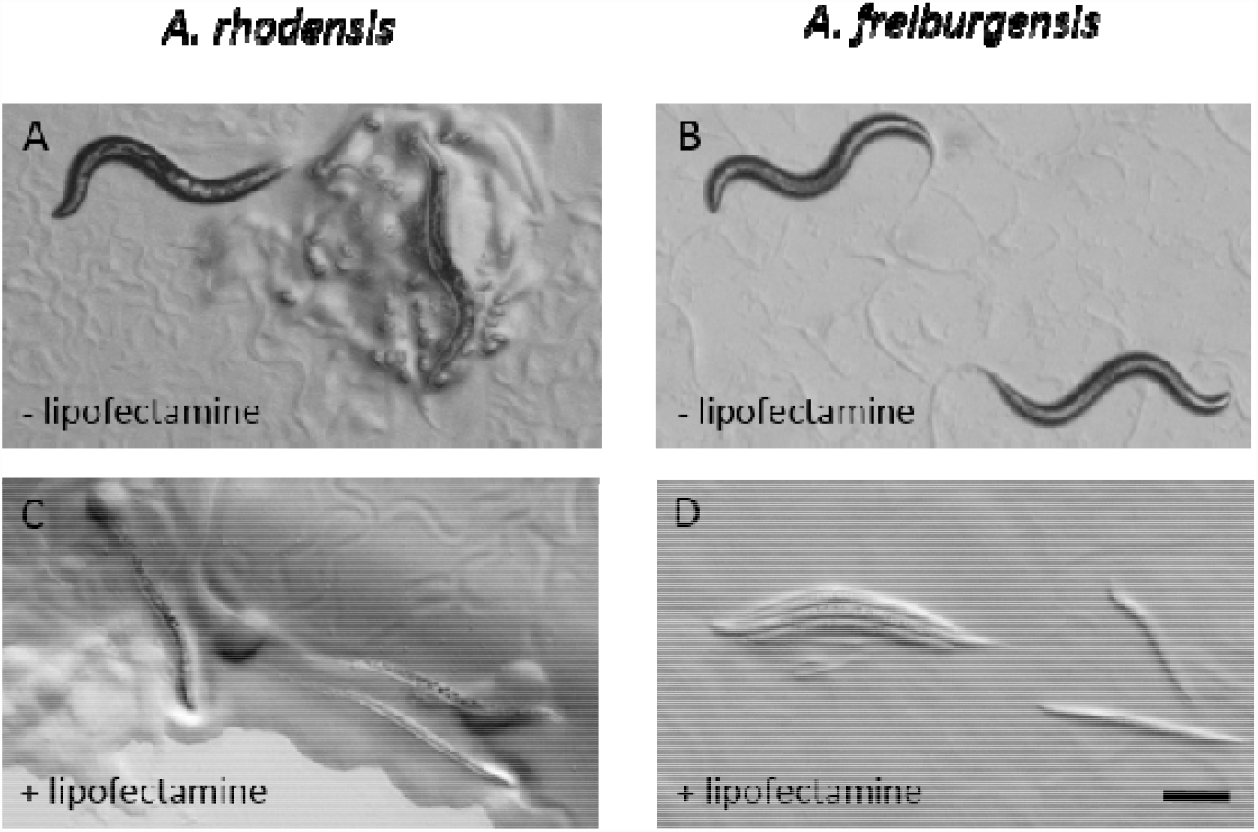
Lipofectamine significantly enhances RNAi against the twitchin homologues, *Arh*-*unc-22* and *Afr-unc-22*, in *A. rhodensis* and *A. freiburgensis*,respectively. Scale bar is 200 µM.

### Lipofection-based methods dramatically improve CRISPR-Cas9 genome mutagenesis in *A. rhodensis* and *A. freiburgensis*

To facilitate screening for CRISPR-Cas9-mediated genome edits, we aimed to introduce a mutation predicted to result in an easily detectable dominant phenotype in the F1 progeny. In *C. elegans* the *rol-6*(*su1006*) allele has been used extensively as a dominant marker, both for transgenesis and for direct genome editing^35,60,61^. *Cel*-*rol-6* encodes a cuticular collagen^62^, and substitution of cysteine for the arginine residue at position 71 results in a dominant right-handed roller phenotype^47^. Furthermore, the same mutation in a similar gene in *P. pacificus* also results in a dominant right-handed roller phenotype^48^. Four potential *rol-6* homologues were identified in *A. rhodensis* and *A. freiburgensis*(Figure S3). The homologues encoding predicted proteins with the highest level of identity to *Cel-* ROL-6, *Arhrol-6.1*, and *Afr-rol-6.1* (62 and 61% identity, respectively), were chosen as target genes for modification (Figure S4). Although the *Cel-rol* homologues in each species exhibit a degree of similarity at the amino acid level (for example, *Arh*-ROL-6.1 and *Arh*-ROL-6.2 share 55% identity), there is significant divergence at the DNA level. Our CRISPR-mediated gene editing strategy uses target-specific sgRNAs and a 100 bp single-stranded DNA donor fragment to introduce a precise gene edit via homologous recombination. The sgRNAs designed are not predicted to anneal to the other *rol-6*-like genes in the respective species, and the DNA donor fragment does not exhibit sufficient homology to act as a repair template in other *rol-6*-like genes (for example, between *Arh-rol-6.1* and *Arh-rol-6.2*, there are 40/100 mismatches in the region of the donor fragment and 9/20 mismatches in the gene-specific crRNA binding region).

The inclusion of lipofectamine dramatically improved the efficiency of the Cas9 mutagenesis at both *Afr-rol-6.1* and *Afr-rol-6.1*(Table 3, Figure 3). Microinjections without lipofectamine did not result in the roller phenotype in the P0 or the F1 progeny of either species. In contrast, if the same complex mixture was introduced with lipofectamine, we observed a distinct roller phenotype in 77% of microinjected *A. rhodensis* P0s and 37% of *A. freiburgensis* P0s, 200 min after injection (n= 27 for both species) (Table 3, Supplemental videos (https://figshare.com/s/e6d12d0b735f7a3f1c09)). Development of the roller phenotype in the P0s could be followed, as the worm tracks changed from a sinusoidal wave to a circular pattern (Figure 3C), presumably as abnormal collagen proteins encoded by the edited gene were incorporated into the cuticle. We observed that, in both species, the P0 worms exhibiting the roller phenotype gave rise to F1 progeny with the same phenotype (Table 3). To verify that roller progeny were produced across the entire lifespan of the injected P0, we phenotyped a subset of *A. rhodensis* injections in detail. P0 individuals produced broods consisting exclusively of roller F1 offspring, whilst non-roller P0s or non-lipofectamine injected controls produced broods consisting of wild-type F1 progeny only (Table 3). Preliminary observations suggested that F1 roller worms went on to produce broods consisting of 100% roller progeny, leading to the assumption that both alleles of the targeted gene were edited. To test this, we tracked inheritance of the phenotype by the F2 progeny from 3 independent lines (Table 3). Each line originated from a single roller P0. In total, the progeny of 22 F1 individuals, representing all 3 lines, were counted and all exhibited the roller phenotype (n= 2,539), indicating that both alleles of the *Ahr-rol-6.1* locus were edited in the F1. The roller phenotype was faithfully inherited, for over 50 generations, in all maintained lines.

**Table 3.** Summary of CRISPR-Cas9 genome mutagenesis in *A. rhodensis, A. freiburgensis* and *S. stercoralis*. Results shown are the combined data from at least 2 independent sets of injections. Hermaphrodite (H), Female (F), dauer larvae (L2d), infective third-stage larvae (iL3). *A. rhodensis* L2d develop into hermaphrodite adults. * Combined total of F2 progeny from 3 independent lines, originating from 3 roller P0 (F1, n= 9, 6 and 7).

| Target gene | Life stage injected | Lip (+/-) | P0 rolling (%) | F1 rolling (%) |  | F2 rolling (%)<br>from rolling F1 parent |
| --- | --- | --- | --- | --- | --- | --- |
|  |  |  |  | P0 rolling | P0 wild-type |  |
| *Arh-rol-6.1* | Adult (H) | Lip + | 77<br>(n= 27) | 100<br>(n= 1144) | 0<br>(n= 466) | 100<br>(n= 2539)* |
|  | Adult (H) | Lip - | 0<br>(n= 13) | NA | 0<br>(n= 1173) | NA |
|  | Larval (L2d) | Lip + | 80<br>(n= 20) | 100<br>(n= 1658) | 0<br>(n= 885) | ND |
|  | Larval (L2d) | Lip- | 0<br>(n= 8) | NA | 100<br>(n= 1958) | ND |
| *Afr-rol-6.1* | Adult (H) | Lip + | 37<br>(n= 27) | 100<br>(n= 1802) | 0<br>(n= 1202) | ND |
|  | Adult (H) | Lip - | 0<br>(n= 10) | NA | 0<br>(n>1000) | ND |
| *Sst- SSTP-0000742500* | Adult (F) | Lip + | 0<br>(n= 15) | NA | 2<br>(n= 200) | NA |
|  | Larval (iL3) | Lip + | 13<br>(n= 23) | 0<br>(n= 5) | ND | NA |

**Figure 3.**
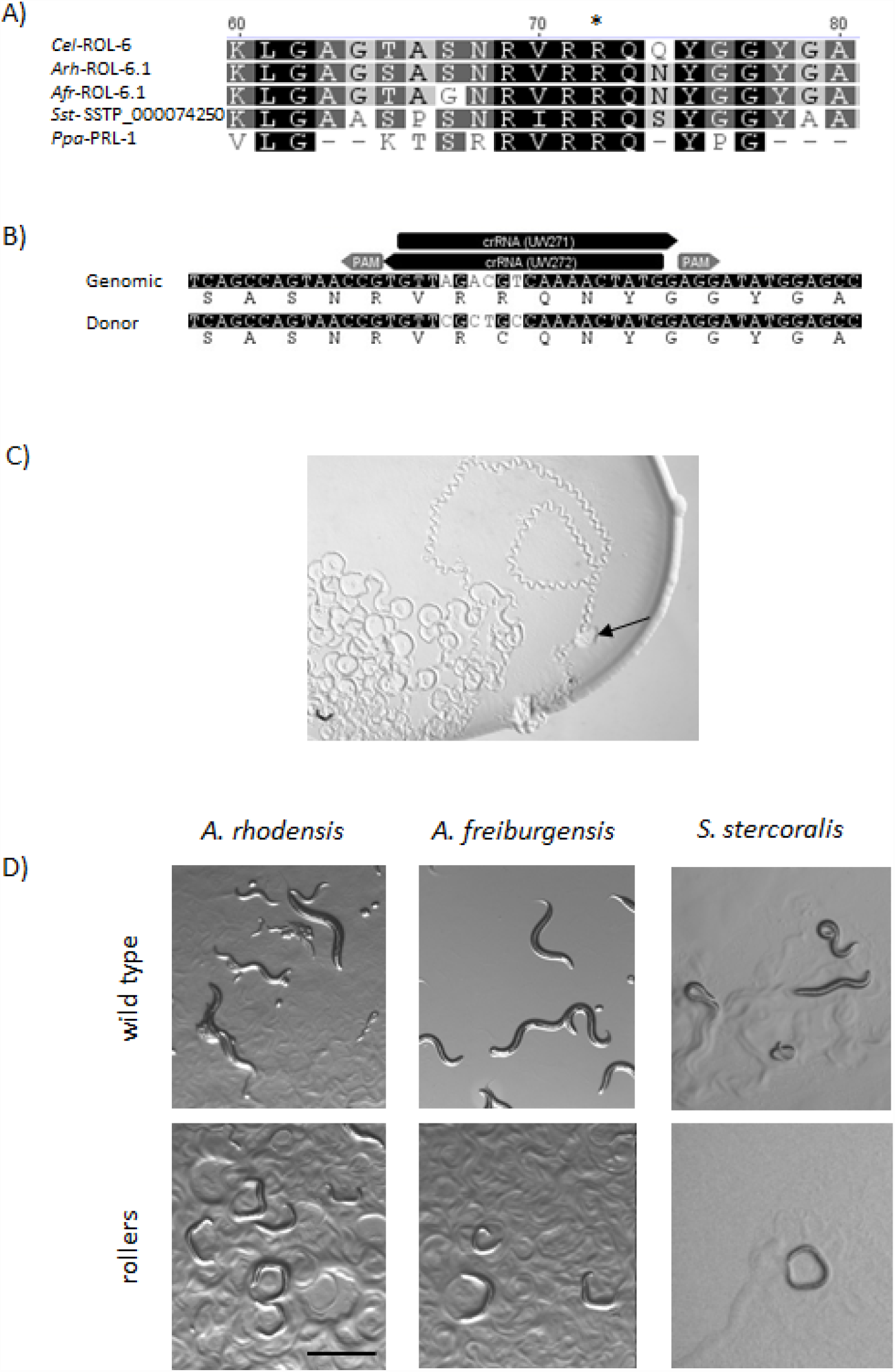
Cas9/CRISPR mediated gene editing of *rol-6* homologues in *A. rhodensis, A. freiburgensis* and *S. stercoralis* results in a right-handed roller phenotype. (A) Alignment of the predicted protein sequences of the ROL-6 homologues encoded by *Cel-rol-6, Arh-rol-6.1*(MH124552), *Afr-rol-6.1* (MH124551), *S. stercoralis* (SSTP_0000742500) and *Ppa-prl-1*. All show conservation of the arginine residue (*) which is converted to cysteine in ROL-6 (*su1006*) in *C. elegans* and *Ppa-* PRL-1 (*tu92*) in *P. pacificus*. (B) Arrangement of the crRNAs, PAM sites and ssDNA donor fragment used for conversion of the *Arh-rol-6.1* gene in *A. rhodensis*. The black arrows indicate the position of gene-specific regions of the anti-parallel crRNA pairs and the grey arrows their corresponding PAM sites. crRNAs represented by forward arrows bind to the non-coding strand, whilst those shown as reverse arrows bind the coding strand. Used in combination, the crRNA antiparallel pairs should result in Cas9-mediated double-stranded DNA breaks on either side of the targeted region. A similar method was used to target *Afr-rol-6.1* and the *S. stercoralis* locus SSTP_0000742500 (Figure S6). (C) Track pattern of *A. rhodensis* adult hermaphrodite 200 min after injection with the gene conversion complexes targeting *Arh-rol-6.1*, as described in (B). Immediately after injection, the individual leaves a wild-type sinusoidal wave as it moves through the bacterial lawn (black arrow highlights the starting point). The pattern becomes increasingly circular as the nematodes movement becomes locked in a clockwise twisting motion, characteristic of the roller phenotype. (D) Right-roller phenotype (bottom panel) compared to wild-type movement (top panel) in *A. rhodensis, A. freiburgensis* (adults and larvae) and *S. stercoralis*(post free-living L2). Scale bar is 1 mm.

Confirmation of the genome changes in stable *Auanema* roller lines proved technically challenging. Although we were able to isolate genomic DNA from roller lines, the DNA was resistant to PCR amplification, either with primer sets designed to amplify the *rol-6.1* locus or those targeting other genomic regions. In contrast, DNA extracted from either *C. elegans* roller lines or wild-type *Auanema*, readily amplified with all primer sets used, suggesting this was a specific issue with the *Auanema* roller lines. Collagen has previously been shown to inhibit PCR^63,64^. Inclusion of a Collagenase I digestion step in the DNA extraction protocol, coupled with the use of a high-fidelity Taq polymerase, allowed PCR amplification across the targeted *Arh*-*rol-6.1* region in a single *A. rhodensis* roller line. Sanger sequencing of this amplicon identified a single base pair change (C to G), resulting in conversion of an arginine to a glycine residue at position 70 of *Arh*-ROL-6.1 (Supplemental Figure S7).

### Lipofection-mediated CRISPR-Cas9 genome editing in free-living stages and infective larvae of the parasitic nematode *S. stercoralis*

One of the main barriers to the genomic manipulation of parasitic nematodes with traditional microinjection techniques is that, with the exception of *Strongyloides* and related parasite genera^65^, the only free-living stages are larvae, with undeveloped gonads. The option to modify different life stages of parasitic nematodes prior to re-infection and to screen for dominant markers could greatly enhance genomic manipulation in such species. Following the simultaneous editing of genes in both the soma and germline of adult *Auanema*, we asked whether we could modify the genome of larval stages of *A. rhodensis* by direct microinjection of the lipofectamine-RNP complexes into the dauer, a developmental stage that resembles the infective third larval stages of many parasitic nematodes. In total, 80% (n= 20) of dauer larvae injected with liposome complexes targeting *Arh-rol-6.1* exhibited the characteristic roller phenotype within 200 mins (Table 3, Supplemental Video (https://figshare.com/s/e6d12d0b735f7a3f1c09)). Although these larvae took a further 24 to 48 h to develop into adults, they went on to produce only roller offspring over their entire reproductive life (1,658 F1 progeny, Table 3). Thus, we were able to simultaneously induce both a screenable dominant phenotype in a larval stage of *A. rhodensis* stemming from a heritable genomic edit, even though the gonad was immature, consisting of only a few cells at the time of treatment. No phenotype was observed if lipofectamine was omitted from the injection mixture in either the injected dauers of *A. rhodensis* or their progeny (1,958 F1 progeny screened from 8 injected L2d larvae, from 2 independent experiments, Table 3).

To ascertain whether the method could be transferred to parasitic species, we aimed to induce a mutation in the SSTP-0000742500 gene of *S. stercoralis*, which encodes a predicted protein with 45% identity to *Cel*-ROL-6 (Figures S3, S4). A sequence-specific donor fragment and crRNA anti-parallel pair were designed to induce an arginine-to-cysteine conversion at the conserved residue in SSTP-0000742500 (Figure 3A, Table S2, Figure S6). Lipofectamine-RNP complexes were injected into the body cavities of infective third-stage larvae (iL3) or the syncytial ovaries of free-living females. Although no injected females exhibited the roller phenotype (n= 48), rollers were observed in a small percentage of their F1 progeny (4%) (Table 3, Supplemental Video (https://figshare.com/s/e6d12d0b735f7a3f1c09). Interestingly, 13% of the iL3 larvae (3/23 surviving P0 nematodes) exhibited the expected roller phenotype 4 to 6 h after microinjection (Table 3, Figure 3D). However, as the small number of roller iL3 generated in this experiment was insufficient to establish a patent infection in susceptible host animals (dogs or gerbils), it was impossible to ascertain whether the CRISPR-induced edit to SSTP-0000742500 in iL3 could be transmitted to the next generation.

### Lipofection facilitates transient expression assays

To develop transgenic protocols in *Auanema*, we produced the species-specific expression vector *PArh*-*eef-1A.1*::TurboRFP::*Arh*-*tbb-2*-3’UTR (pAS19). Since the genome assembly for *A. freiburgensis* is not currently available and attempts to produce *A. freiburgensis* lines expressing *PArh*-*eef-1A.1*::TurboRFP::*Arh*-*tbb-2*-3’UTR were unsuccessful (no F1 expressing progeny from 40 injected P0), this work focused on *A. rhodensis*.

In preliminary tests, pAS19 was introduced into the gonads of young adult hermaphrodites of *A. rhodensis* by conventional microinjection methods. However, no RFP expression was detected in the F1 progeny from over 100 injected individuals (data not shown). In contrast, co-microinjection of pAS19 with lipofectamine resulted in 20% of injected P0 (n= 10), producing F1 progeny expressing RFP (16 F1 expressing RFP, from 210 progeny, from 2 independent injections). In *C. elegans, eef-1A.1* encodes a translation elongation factor that is required for germline maintenance and embryonic viability^66–68^. *Cel* - *eef-1A.1* is expressed in the soma throughout development as well as in the adult germline (Mitrovich, 2000 #14009; Frøkjaer-Jensen, 2012 #14080}. However, in *A. rhodensis PArh*-*eef-1A.1*::TurboRFP the expression was primarily limited to embryonic development (Figure 4). No RFP expression was observed in the F2 progeny from 16 F1 lines (over 200 progeny screened per line), and the RFP transgene was not detected by PCR in any of the genotyped F2 individuals (n= 10, data not shown).

**Figure 4.**
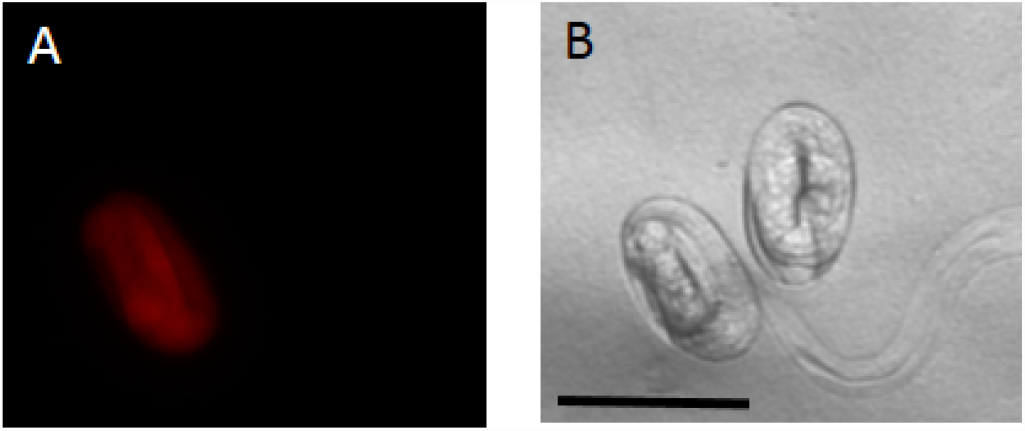
Microinjection of liposome-DNA complexes facilitates expression of the P*eef-1A.1*::TurboRFP::*tbb-2*-3’UTR transgene in *A. rhodensis*. Expression was primarily limited to late embryogenesis in F1 progeny. Scale bar 50 µM.

## Discussion

The success of functional genomic studies in non-model nematodes has been variable, and many species seem to resist different methods^21^. By using liposome-based transfection reagents, we were able to establish RNAi and CRISPR-Cas9 mutagenesis, as well as transient transgene expression in nematodes with cellularised germlines, such as those present in *Auanema* nematodes. Liposomes mediate the transfer of the dsRNA and CRISPR-Cas9 components through cells from the point of injection. They can then spread to the rest of the gonad and to the rest of the body of the injected animal. Injected animals frequently showed the expected RNAi-or CRISPR-Cas9-induced phenotype within 24 hours after injection. The progeny of these injected animals inherited the phenotype affecting, in many cases, 100% of the F1 progeny.

Although RNAi and CRISPR-Cas9 mutagenesis are reliable technologies in *Auanema*, establishing a robust system for transgene expression has been more challenging. There could be many reasons for the lack of *PArh-eef-1A.1*::TurboRFP transgene expression in the germline of *A. rhodensis*, which was initially expected based on the homology of *Arh-eef-1A.1* to *Cel-eef-1*. For example, *A. rhodensis Arh-eef-1A.1* may only be expressed during embryonic development. Alternatively, there may be technical issues associated with the transgene. For example, the transgene may be silenced during development, may not be transmitted to the germline tissue, or the transgene may be suboptimal for germline expression. Indeed, in the individuals we tested, the transgene did not pass from the F1 to the F2 progeny (data not shown). Future work will aim to produce more stable transgenic lines, using methods as implemented in *P. pacificus*, to induce genomic integration of the transgene^48^. Further optimisation of germline expression of transgenes in *Auanema* will be important. In *C. elegans*, for example, the choice for the 3’ UTR is critical for driving germline expression^69^. For instance, *eef-1A.1* promoter-driven germline expression of a transgene was observed in *C. elegans* only when paired with the *tbb-2* 3’UTR, but not with the *unc-54* 3’ UTR^68,70^.

The addition of lipofectamine to the injection mixture not only helped in establishing RNAi in *Auanema* but also in *P. pacificus*. *P. pacificus* is a nematode model used for evolutionary studies^71^, for which successful RNAi has been restricted to the silencing of a gene conferring gain-of-function phenotypes^32,34^. Although *P. pacificus*, like *C. elegans*, has a syncytial germline^72^, this part of the gonad is much smaller and difficult to inject because of the gonad topology. Therefore, the gain in efficiency on inclusion of lipofectamine may be due to a better spread of the injection mixture throughout the germline.

Our results indicate that injection of CRISPR-Cas9 components with lipofectamine allowed efficient mutagenesis of the *Auanema* genome for the first time. In almost all individuals injected with *Arh-rol-6.1* CRISPR-Cas9, the entire F1 broods had the expected right-handed roller phenotype. The fact that the injected animals generated F1s that segregated the roller phenotype in 100% of the F2s indicates that the F1s are homozygous for the mutation. We identified a base pair substitution in *Arh-rol-6.1* line 5 which results in conversion of a second conserved arginine adjacent to the targeted residue.

We also succeeded with CRISPR-Cas9 mutagenesis in *S. stercoralis*, a nematode parasite of humans and dogs. The fact that the host-dwelling life stages are inaccessible is one of the many challenges of inducing heritable phenotypes in parasitic nematodes^73^ because it is difficult to deliver the CRISPR-Cas9 machinery to the germlines of the reproductive stages. *Strongyloides* sp., however, is one of the few parasitic nematodes in which genome modifications have been successful^74,75^. Adult germlines of *Strongyloides* spp. are accessible because they have a reproducing free-living generation. With the addition of lipofectamine to the CRISPR-Cas9 injection mixture, we were able to induce a transmissible phenotype from the injected free-living females to larvae of the F1 generation. Remarkably, we also observed the expected phenotype in iL3 directly injected with CRISPR-Cas9 elements with lipofectamine. Thus, this technique could be used in other parasitic nematodes species that lack a reproducing free-living generation by inducing mutations in the germlines of pre-infective or infective larval stages that would be retained as they developed to parasitic stage in the host.

While the roller phenotypes conferred by dominant mutations in *rol-6* homologues constitute useful markers for transgenesis or gene editing in free-living nematodes, their utility in parasitic nematodes is doubtful. The low numbers of roller *S. stercoralis* iL3 produced in this study did not permit an attempt at host passage to ascertain the infectivity of those roller larvae and the heritability of this mutation and its associated phenotype. Additionally, considering the fast, progressive motility of iL3 of *S. stercoralis* and many other parasitic nematodes at the time of host infection, we consider it likely that the impaired progressive motility of roller mutants would compromise or completely negate their ability to infect and complete tissue migrations required to enter and persist within their predilection sites in the host. Nevertheless, this dominant phenotype, with its rapid onset and ease of recognition, allowed proof of principle for the lipofectamine-facilitated transfer of CRISPR elements to somatic cells of *S. stercoralis* iL3 and provides a strong basis for further study of this method. Firstly, it will be important to ascertain heritability of lipofectamine-facilitated genome modifications in *Strongyloides* iL3 or other pre-infective larval stages. This would require a host passage, so to avoid adverse effects of motility phenotypes on infectivity, we submit that this should be addressed by deploying lipofectamine technology to facilitate the CRISPR-mediated integration of a fluorescent reporter transgene into a non-coding region of the *Strongyloides* genome under study during the iL3 or another preinfective larval stage. Extrapolating from the present results, expression of such a reporter should ensue within hours of injection, allowing selection of fluorescent individuals for host passage. Given the availability of a natural rodent host, the rat, that allows successful passage of small numbers (10–20) of infective larvae, *Strongyloides ratti* would be the most practical subject for such an experiment. Secondly, as alluded to above, it will be crucial to obtain proof of principle for this mutagenesis strategy in parasitic nematode species that have free-living pre-infective and infective larvae but lack an entire free-living generation. This description applies to soil-transmitted nematodes of paramount medical and agricultural importance, the hookworms, and trichostrongyles. The logical subjects for a pilot study of genome editing in such parasites could be their counterparts infecting laboratory rodents, *Nippostrongylus brasiliensis* and *Heligmosomoides polygyrus*,respectively.

In summary, the addition of an inexpensive liposome-based transfection reagent has opened the door for exciting future functional genomic studies in non-model nematode systems, including both free-living and parasitic species.

## Acknowledgements

P.P. was funded by University of Warwick start-up funds. S.A. and A.P.-d.S. were funded by BBSRC (BB/L019884/1) and University of Warwick start-up funds. J.B.L. acknowledges support from grants AI50688, AI105856 and OD P40–10939 from the US National Institutes of Health.

## Author Contributions Statement

S.A. and A.P.-d.S. are responsible for the idea, concept, and design of the experiments. S.A. and P.P. conducted the experiments in *Auanema*. The experiments in *S*. *stercoralis* were conducted by H.S. under the supervision of J.B.L. S.A., J.B.L. and A.P.-d.S. wrote the final manuscript with the assistance of the co-authors. All authors reviewed the manuscript.

## Competing financial interests

The authors declare no competing financial interests.

